# Changes in nucleus accumbens core translatome accompanying incubation of cocaine craving

**DOI:** 10.1101/2024.09.15.613147

**Authors:** Alex B. Kawa, Joel G. Hashimoto, Madelyn M. Beutler, Marina Guizzetti, Marina E. Wolf

## Abstract

In the ‘incubation of cocaine craving’ model of relapse, rats exhibit progressive intensification (incubation) of cue-induced craving over several weeks of forced abstinence from cocaine self-administration. The expression of incubated craving depends on plasticity of excitatory synaptic transmission in nucleus accumbens core (NAcC) medium spiny neurons (MSN). Previously, we found that the maintenance of this plasticity and the expression of incubation depends on ongoing protein translation, and the regulation of translation is altered after incubation of cocaine craving. Here we used male and female rats that express Cre recombinase in either dopamine D1 receptor- or adenosine 2a (A2a) receptor-expressing MSN to express a GFP-tagged ribosomal protein in a cell-type specific manner, enabling us to use Translating Ribosome Affinity Purification (TRAP) to isolate actively translating mRNAs from both MSN subtypes for analysis by RNA-seq. We compared rats that self-administered saline or cocaine. Saline rats were assessed on abstinence day (AD) 1, while cocaine rats were assessed on AD1 or AD40-50. For both D1-MSN and A2a-MSN, there were few differentially translated genes between saline and cocaine AD1 groups. In contrast, pronounced differences in the translatome were observed between cocaine rats on AD1 and AD40-50, and this was far more robust in D1-MSN. Notably, all comparisons revealed sex differences in translating mRNAs. Sequencing results were validated by qRT-PCR for several genes of interest. This study, the first to combine TRAP-seq, transgenic rats, and a cocaine self-administration paradigm, identifies translating mRNAs linked to incubation of cocaine craving in D1-MSN and A2a-MSN of the NAcC.

## Introduction

One defining characteristic of substance use disorder is difficulty remaining abstinent. An important contributor to relapse vulnerability is exposure to drug-associated cues that provoke craving. Such cue-induced craving progressively increases, or intensifies, during a drug-free period. This is termed incubation of craving. Incubation occurs in humans across several drug classes [1–4], and can be modeled in rodents following cocaine use [5–8].

The nucleus accumbens (NAc) is critical for incubation of cocaine craving (see below). It is composed primarily of medium spiny neurons (MSN) that integrate excitatory inputs from cortical and limbic regions and project to downstream regions of the basal ganglia to control motivated behavior. MSN of the NAc express either dopamine type-1 receptors (D1-MSN) or dopamine type-2 receptors (D2-MSN). Generally, D1-MSN are involved in initiating and driving motivation, whereas D2-MSN have been found to be less involved in motivated behavior or oppose it [9–11]. Current theories emphasize the importance of D1-MSN/D2-MSN pathway balance in shaping behavior [12–15].

Incubation of craving during abstinence from cocaine self-administration ultimately depends on strengthening of excitatory synapses on NAc core (NAcC) and shell (NAcS) MSN through upregulation of Ca^2+^-permeable AMPARs (CP-AMPARs) [16,17], although the underlying cascade may differ between the two subregions [18–22]. In NAcC, CP-AMPAR elevation occurs after ∼1 month of abstinence from extended-access cocaine self-administration and then persists unabated through at least abstinence day (AD) 70 [23,24]. CP-AMPAR blockade or their removal from synapses prevents incubated cocaine seeking [23–26].

Incubation of cocaine craving and CP-AMPAR upregulation is associated with dysregulation of protein translation in NAcC on multiple levels [27,28], and this dysregulation supports incubation. For example, disrupting protein translation in NAcC of cocaine-incubated rats normalizes CP-AMPAR levels [29] and prevents expression of incubation [28]; these results are summarized in Supplementary Fig. S1. We previously showed that translation of the GluA1 subunit is increased after incubation [27], but other changes in translation remain unknown. To address this gap, we used translating ribosome affinity purification (TRAP) [30–35] to isolate actively translating mRNAs from either D1-MSN or D2-MSN at two timepoints after cocaine or saline self-administration in rats, followed by RNA sequencing. This TRAP-seq approach revealed very few differentially translated (DT) genes on AD1 between rats that self-administered saline vs. cocaine, in either D1-MSN or D2-MSN. However, in cocaine rats, there were hundreds of DT genes on AD40-50 compared to AD1, with substantially more differential translation occurring in D1-MSN. Our study, the first to use TRAP-Seq to distinguish D1-MSN and D2-MSN after cocaine self-administration, provides a unique window into protein translation occurring in NAcC before and after incubation of cocaine craving.

## Materials and Methods

Extended Materials and Methods are provided in Supplementary Materials.

### Subjects and surgery

Procedures were approved by the OHSU IACUC in accordance with the U.S. PHS Guide for the Care and Use of Laboratory Animals. Adult male (n=35) and female (n=31) Long-Evans *Drd1(iCre)* (D1:Cre) and *Adora2a(iCre)* (A2a:Cre) transgenic rats [36] underwent long-access cocaine or saline self-administration (6 h/day × 10 days). A2a is a marker for D2-MSN. These rats received jugular catheter implantation and intracranial surgery to infuse TRAP virus (EF1a-AAV5-FLEX-GFPL10a; Addgene #98747-AAV5) into NAcC. The TRAP virus has been used in mice [33,35], but this represents its first use to study addiction-related plasticity in rats.

### TRAP-Seq

TRAP was performed on NAcC tissue as previously described [37–39]. Six groups were generated for sequencing: D1_AD1 Saline (Sal), D1_AD1 Cocaine (Coc), A2a_AD1 Sal, A2a_AD1 Coc, D1_AD40 Coc, and A2a_AD40 Coc. AD1 rats were taken from home cages and euthanized ∼18 h after the final self-administration session. Rats designated ‘AD40’ were taken from home cages and killed on AD40-50, but are referred to as AD40 for simplicity. Each group consisted of 8 samples (1 sample = bilateral NAcC from 1 rat), split equally between the sexes except for the A2a_AD1 Sal group (5 male/3 female). The significance of RNA-Seq data was determined using DESeq2 FDR-adjusted Wald test with adjusted p-values of <0.05. All processed and raw sequencing reads are publicly accessible at the Gene Expression Omnibus (GEO; https://www.ncbi.nlm.nih.gov/geo/) using accession number GSE263646.

### Bioinformatics analysis

Gene Ontology (GO) and pathway enrichment analysis were carried out using the web-based Panther [40,41] and web-based enrichR package [42,43].

## Results

### Cocaine self-administration

Our goal was to compare translating mRNAs in D1-MSN and A2a/D2-MSN (hereafter referred to as A2a-MSN) of NAcC during incubation of cocaine craving, taking advantage of D1:Cre and A2a:Cre rat lines to enable cell type-specific expression of AAV-TRAP (Fig. 1A, timeline; Fig. 1C, schematic of workflow). Fig. 1B shows self-administration data for the 32 cocaine rats ultimately used for RNA-sequencing, while saline self-administration data are shown in Fig. S2. D1:Cre and A2a:Cre cocaine rats did not differ in number of infusions [genotype: F(1,47)=0.476, p=0.49]. Infusions increased across sessions (‘escalation’) [session: F(9,138)=4.385, p<0.001], and there was no session X genotype interaction [F(9,138)=0.609, p=0.79]. After self-administration, rats were euthanized on AD1 or AD40-50 (referred to as AD40) and actively translating mRNA was isolated from either D1-MSN or A2a-MSN of NAcC. Thus, six groups were generated (D1_AD1 Sal, D1_AD1 Coc, A2a_AD1 Sal, A2a_AD1 Coc, D1_AD40 Coc, A2a_AD40 Coc; n=8 rats/group).

**Fig. 1.**
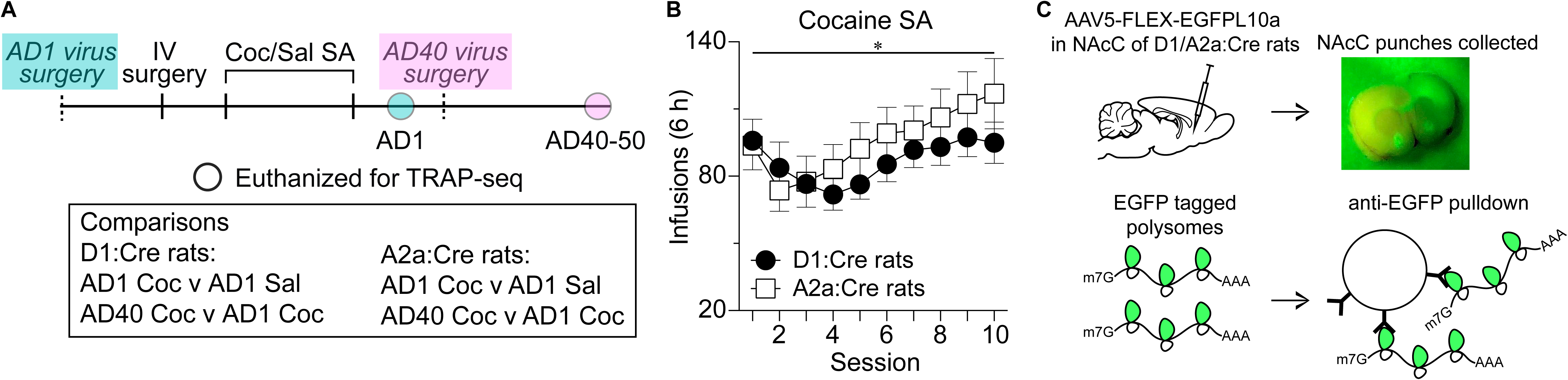
D1:Cre and A2a:Cre rats were infused with AAV5-FLEX-EGFPL10a in NAc core (NAcC) and self-administered cocaine or saline before NAcC tissue was collected at various timepoints and translating mRNA was extracted for sequencing. **A)** Experimental timeline. All rats underwent 10 days of intravenous cocaine or saline self-administration (6 h/day x 10 days). To achieve equivalent virus expression time, rats in the abstinence day (AD) 1 groups had virus infused ∼2 weeks before jugular catheter surgery, while rats in the AD40 groups had virus infused in the week following self-administration. Rats were euthanized for TRAP-seq on either AD1 or AD40-50 (referred to as AD40). There were 6 experimental groups: D1_AD1 Sal, D1_AD1 Coc, D1_AD40 Coc, A2a_AD1 Sal, A2a_AD1 Coc, and A2a_AD40 Coc. **B)** Cocaine self-administration data over 10 sessions in D1:Cre (n=16) and A2a:Cre (n=16) rats. Pokes in the active hole resulted in an infusion of cocaine (0.5 mg/kg) paired with a light cue. D1:Cre and A2a:Cre rats did not differ in the number of infusions self-administered. * Main effect of session on infusions (*F*9,138 =4.385, *p*<.001) with post hoc contrast showing more infusions on sessions 8–10 than sessions 1–3 (paired *t*-test with Bonferroni correction, *p*<0.001). **C)** Schematic depicting tagging and isolation of translating mRNAs. Punches of the NAcC were collected on either AD1 or AD40 following self-administration training. Next, the EGFP tagged polysomes (ribosomes and actively translated mRNA) were immunoprecipitated using protein A/G coated magnetic beads and EGFP antibodies to obtain actively translating mRNAs from D1-MSN or A2a-MSN. RNA was then isolated and sequenced (see Methods for details). AD, abstinence day; IV surgery, intravenous catheter implantation; Coc, cocaine; Sal, saline; NAcC, nucleus accumbens core. Data from the rats that self-administered saline can be found in Supplementary Fig. S2.

### Cre expression in D1:Cre and A2a:Cre rats

Before sequencing samples from cocaine or saline rats, it was necessary to confirm that Cre expression was limited to D1-MSN in D1:Cre rats and A2a-MSN in A2a:Cre rats. In drug-naïve rats (n=3 D1:Cre and 3 A2a:Cre), we used RNAscope to identify *Drd1*, *Drd2*, *Adora2a* (A2a), and *iCre* mRNA. Representative images are shown in Fig. 2A. Visual inspection revealed no overlap between *Drd1* and *iCre* in the A2a:Cre rats, no overlap between *Adora2a* or *Drd2* and *iCre* in the D1:Cre rats, and nearly complete overlap between *Drd1* and *iCre* in the D1:Cre rats and *Adora2a* or *Drd2* in the A2a:Cre rats.

**Fig. 2.**
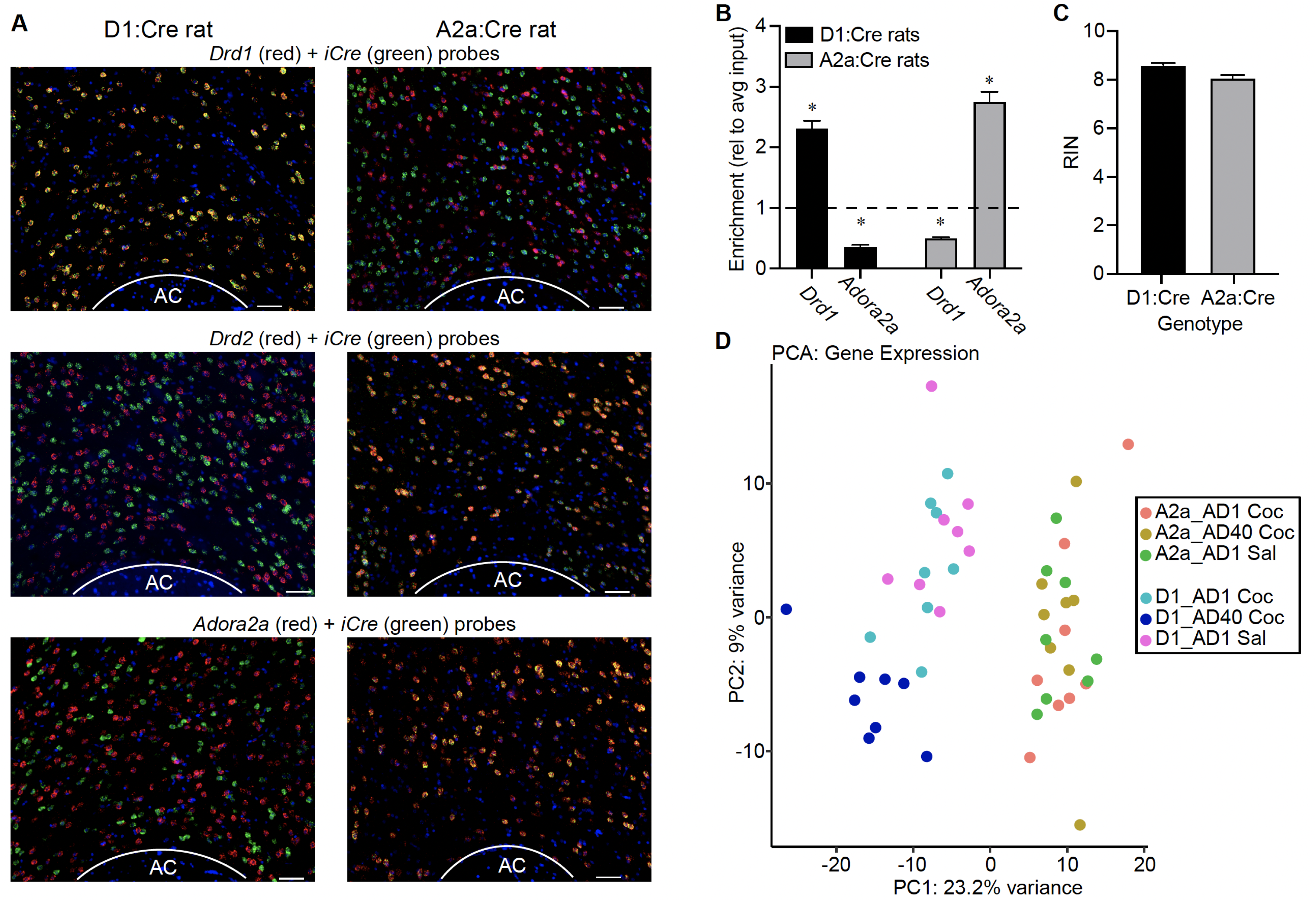
Validation of selective Cre expression in D1-MSN and A2a-MSN of D1:Cre and A2a:Cre rats. **A)** Representative images of fluorescent in situ hybridization in tissue obtained from D1:Cre (left column) and A2a:Cre (right column) rats and treated with a *Drd1* probe (red) and an *iCre* probe (green) (top row), a *Drd2* probe (red) and an *iCre* probe (green) (middle row), and an *Adora2a* probe (red) and an *iCre* probe (green) (bottom row). Overlap of two probes is shown in yellow. Nuclei are stained with DAPI (blue). There was no overlap between *Cre* expression and *Drd2* mRNA or *Adora2a* mRNA in the D1:Cre rats and similarly no overlap between *Cre* expression and *Drd1* mRNA in the A2a:Cre rats. There was near complete overlap between *Cre* expression and *Drd1* mRNA in the D1:Cre rats and near complete overlap between *Cre* expression and *Drd2* mRNA and *Adora2a* mRNA in the A2a:Cre rats. (Scale bars=50 μm). **B)** RT-PCR data from translating mRNA obtained from D1:Cre (n=24) and A2a:Cre (n=24) samples that were later sequenced. Relative to homogenate (input) mRNA, the mRNA from D1:Cre rats was enriched for *Drd1* mRNA and de-enriched for *Adora2a* mRNA, and the mRNA from A2a:Cre rats was enriched for *Adora2a* mRNA and de-enriched for *Drd1* mRNA (*p<0.001, paired *t*-test between input mRNA and TRAP pulldown mRNA of D1:Cre or A2a:Cre rats). **C)** Mean RNA Integrity Number (RIN) of TRAP samples from D1:Cre and A2a:Cre rats that were sequenced. **D)** Principal component analysis showing all 48 samples that were sequenced, color coded by group. PC1 can be identified as cell type with D1 samples clustering to the left and A2a samples clustering to the right. PC2 differentiates D1_AD40 Coc samples from D1_AD1 Coc and D1_AD1 Sal samples, while not differentiating the A2a samples. AC, anterior commissure

The TRAP pulldown samples that were eventually sequenced were also tested for Cre recombination selectively in D1-MSN or A2a-MSN using qRT-PCR to quantify *Drd1* and *Adora2a* mRNA. As expected, in the D1-TRAP samples *Drd1* mRNA was enriched (p<0.001) and *Adora2a* mRNA was de-enriched (p<0.001) (compared to input fraction: bulk NAc homogenate before TRAP immunoprecipitation), while *Drd1* mRNA was de-enriched (p<0.001) and *Adora2a* mRNA was enriched (p<0.001) in the A2a-TRAP samples (Fig. 2B). We further validated our TRAP protocol and fidelity of Cre expression in drug-naïve rats by showing that the D1-MSN markers *Drd1* and *Pdyn* were enriched in D1-TRAP samples and depleted in A2a-TRAP samples, *Adora2a* and *Drd2* were enriched in A2a-TRAP samples and depleted in D1-TRAP samples, and the astrocyte marker *Gfap* is depleted in both D1- and A2a-TRAP samples (Fig. S3).

### Cell-type differences in sequencing results

Prior to sequencing, the TRAP pulldown mRNA underwent quality control via Bioanalyzer. The mean (±SEM) RNA Integrity Numbers (RIN) for D1-TRAP and A2a-TRAP samples were 8.61±0.08 and 8.08±0.11, respectively (Fig. 2C). No samples with RIN <7 were sequenced. As expected from prior studies comparing D1-MSN and A2a/D2-MSN transcriptomes [34,44–46], principal component (PC) analysis (Fig. 2D) revealed that PC1 (23.2% of variance) mapped onto cell type (D1 vs. A2a). Visually, D1-TRAP samples cluster to the left, and A2a-TRAP samples cluster to the right.

### Cocaine self-administration and an abstinence period change the translatome in D1-MSN

Interestingly, PC2 (9% of variance) mapped onto AD for D1-TRAP samples (AD1 vs. AD40), while this was not apparent for A2a-TRAP samples (Fig. 2D). Visual inspection of the PCA plot in Fig. 2D reveals that the D1_AD40 Coc samples cluster together in the lower left of the plot, and the D1_AD1 Coc samples and D1_AD1 Sal samples cluster in the upper left. This pattern is not observed for A2a samples.

Planned contrasts between D1_AD1 Coc and D1_AD1 Sal samples showed that there were 0 differentially translated (DT) genes between these groups (all genes p-adj.>0.05; Fig. 3A). The same comparison run on A2a-TRAP samples (AD1_Coc vs. AD1_Sal) showed 2 genes that were upregulated in the AD1_Coc samples, relative to AD1_Sal samples, and no downregulated genes (Fig. 3B). In contrast, the comparison between D1_AD40 Coc vs. D1_AD1 Coc samples revealed 521 DT genes, indicating that incubation of craving is accompanied by changes in the D1-MSN translatome (216 up; 305 down; p-adj.<0.05; Fig. 3C). The same comparison in the A2a-TRAP samples (AD40_Coc vs. AD1_Coc) showed 71 DT genes (25 up; 46 down; p-adj.<0.05; Fig. 3D). These contrasts are summarized in Fig. 3E for D1-TRAP samples (top) and A2a-TRAP samples (bottom).

**Fig. 3.**
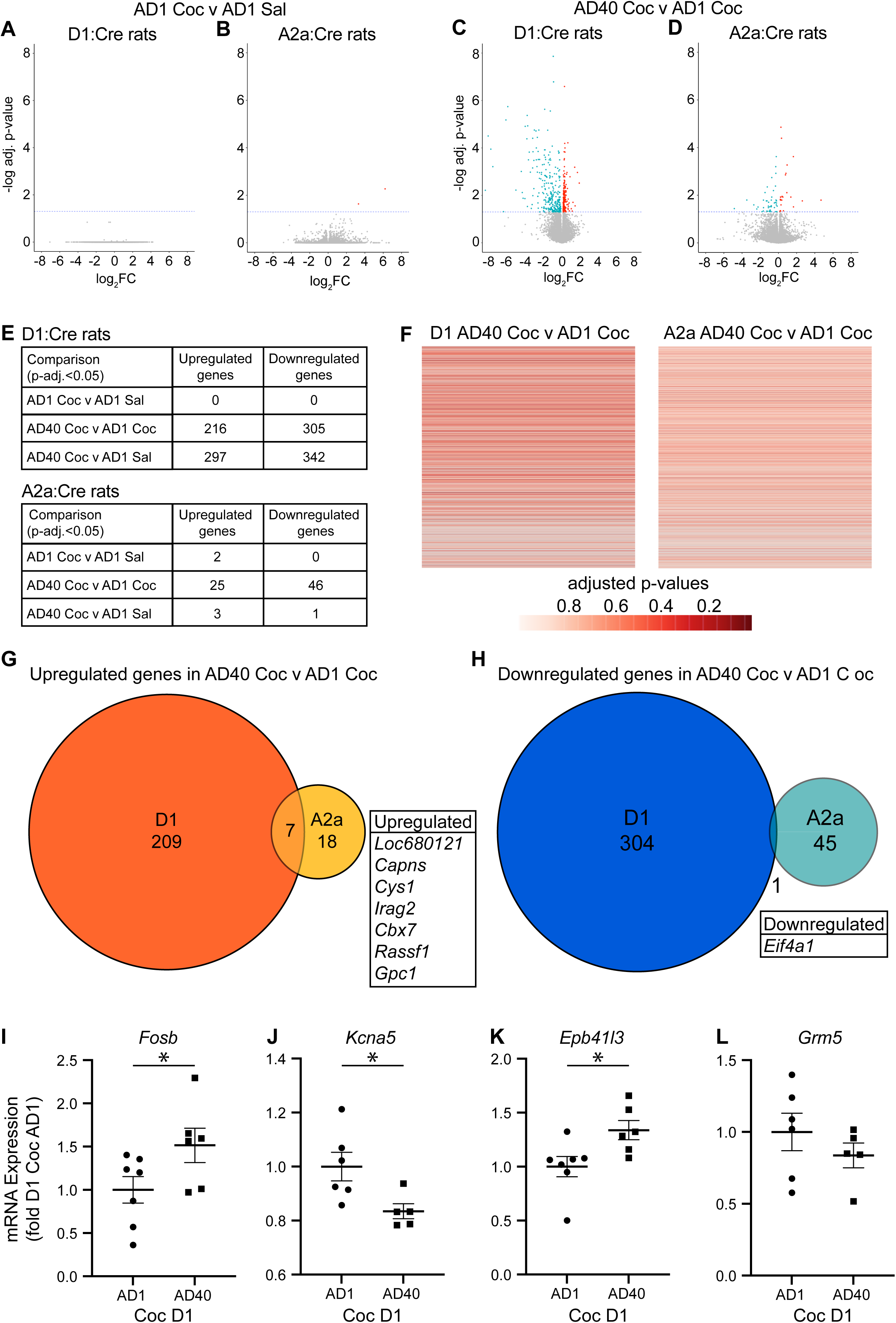
Sequencing data from D1:Cre and A2a:Cre TRAP samples. **A-D)** Volcano plots showing the adjusted p-value and Fold Change (FC) for genes from the AD1 Coc vs. AD1 Sal comparison (A&B) and AD40 Coc vs. AD1 Coc comparison (C&D) for D1:Cre rats and A2a:Cre rats. Red dots represent significantly upregulated genes and blue dots represent significantly downregulated genes (adj. p-value<0.05). **E)** Summary tables depicting the number of differentially translated (DT) genes for each comparison listed in the table. Notably there were very few DT genes following cocaine self-administration when rats were euthanized the day after the last session (AD1), compared to saline self-administering rats. There were far more DT genes when AD40 Coc and AD1 Coc groups were compared, and this contrast was more robust in the D1:Cre rats than A2a:Cre rats. **F)** Heat map showing the results of the D1_AD40 Coc vs. D1_AD1 Coc comparison (left) and the A2a_AD40 Coc vs. A2a_AD1 Coc comparison (right). Each row represents a gene. The rows/genes are aligned in the two heat maps. ‘Hotter’/darker red colors indicate a lower adjusted p-value. **G-H)** Venn diagrams showing the overlap of upregulated genes (G) and downregulated genes (H) between D1:Cre and A2a:Cre rats for the AD40 Coc vs. AD1 Coc comparisons. The genes that were upregulated or downregulated in both D1: and A2a:Cre rats are listed in the inset tables. N = 8 rats/group (4M, 4F) for sequencing analysis. **I-L)** qRT-PCR validation of genes of interest identified as differentially translated in D1 MSN by TRAP-Seq. Comparing AD40 to AD1 for D1-MSN, TRAP-seq analysis indicated differential translation (p-adj.<0.05) of *Fosb* (upregulated), *Kcna5* (downregulated), *Epb41l3* (upregulated), and *Grm5* (downregulated). The potential relationship of these genes to incubation of cocaine craving is considered in the Discussion. qRT-PCR was performed on aliquots of the same samples used for RNA-seq, except for 3 samples (1 Male AD1, 1 Male AD40, 1 Female AD40) for which insufficient quantities of RNA remained after RNA-seq. Methods and primers are detailed in Supplementary Methods. qRT-PCR confirmed the significant upregulation of *Fosb* (p = 0.0308) and *Epb41l3* (p = 0.0127), and downregulation of regulation of *Kcna5* (p = 0.0142). *Grm5* showed the same direction of regulation in qRT-PCR analysis as seen TRAP-seq analysis but was not significant in the former (p = 0.1715). A gene was “upregulated” (or “downregulated”) if it was increased (or decreased) in the first group listed, relative to the second group. N=6-7 rats/group (D1 MSN) and 5-6 rats/group (D2 MSN) for qRT-PCR.

Fig. 3F shows heat maps comparing AD40 Coc vs. AD1 Coc for D1-TRAP and A2a-TRAP samples. Each gene is represented by a row, with ‘hotter’/darker red colors indicating lower p-values in the AD40 Coc vs. AD1 Coc comparison. The heat maps illustrate more robust changes over 40 days of abstinence from cocaine self-administration in D1-TRAP samples (left) than in A2a-TRAP samples (right).

Of the 216 genes upregulated in the D1_AD40 Coc vs. D1_AD1 Coc comparison, only 7 were also upregulated in the A2a_AD40 Coc vs. A2a_AD1 Coc comparison (Fig. 3G; listed in table inset). Of the 305 genes downregulated in the D1_AD40 Coc vs. D1_AD1 Coc comparison, only 1 was also downregulated in the A2a_AD40 Coc vs. A2a_AD1 Coc comparison (Fig. 3H; listed in table inset).

Overall, these results suggest that there is very little translational reprogramming induced by cocaine vs. saline self-administration when translating mRNA is measured ∼24 hours after the last self-administration session and no stimulus (e.g. cue, cocaine, context) is presented prior to tissue collection. However, there were far more translational changes after incubation plateaued (when compared to AD1 Coc or AD1 Sal samples) and these changes were present in both D1-MSN and A2a-MSN, but to a much greater extent in D1-MSN.

To confirm the sequencing results, the sequenced samples were further assessed using qRT-PCR for select genes of interest (Fig. 3I-L). The sequencing results showed that in the D1_AD40 Coc vs. D1_AD1 Coc comparison, translation of *Fosb* and *Epb41l3* was upregulated on AD40 while downregulation was observed for *Kcna5* and *Grm5* (p-adj.<0.05). The qRT-PCR results confirmed the sequencing findings for *Fosb* (one-tailed *t*-test; p=0.031), *Kcna5* (p=0.014), *Epb41l3* (p=0.013), and *Grm5* showed a change in the same direction found with sequencing (p=0.172). For more information on these genes see Discussion.

### Sex differences in the translatome response in D1-MSN and A2a-MSN

Fig. 4A shows the same PCA plot depicted in Fig. 2D, but in this case the points corresponding to individual samples are color-coded by sex. This PCA plot does not indicate any clustering by sex.

**Fig. 4.**
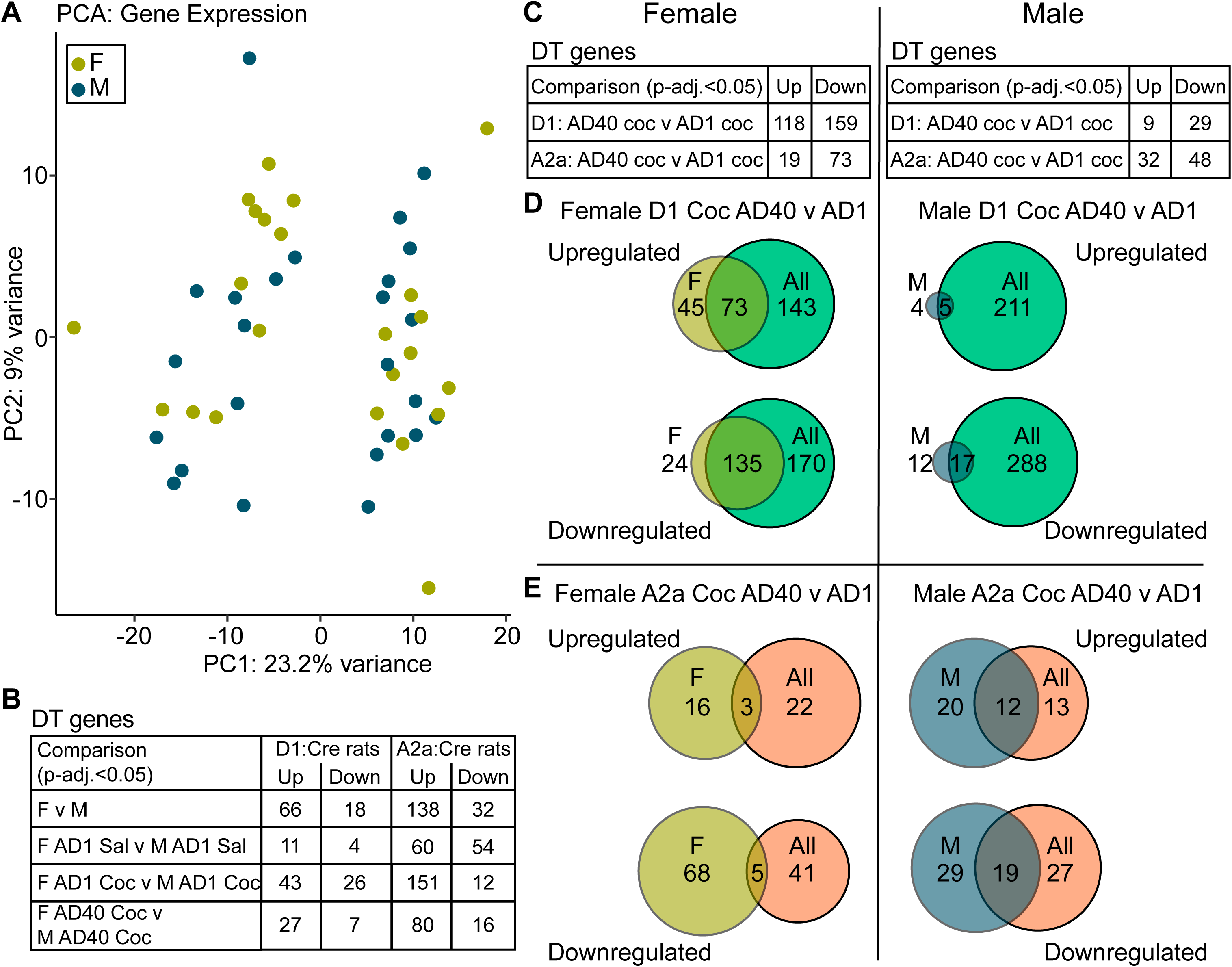
Comparison of sequencing data for male and female rats. **A)** Principal component analysis plot color-coded by sex (the same plot is shown color-coded by group in Fig. 2). There is no obvious clustering of sex along PC1 or PC2. **B)** Table summarizing the results of specified comparisons *between* the sexes for differentially translated (DT) genes. For each comparison shown, there were more DT genes (sex-dependent regulation) in the A2a:Cre samples than the D1:Cre samples. **C)** Tables summarizing the results of the specified comparisons *within* each sex. There were substantially more DT genes in females (left) than males (right) when comparing AD40 Coc vs. AD1 Coc in D1:Cre samples, but not in A2a:Cre samples. **D-E)** Venn diagrams showing the Coc AD40 vs. Coc AD1 comparison in D1:Cre **(D)** and A2a:Cre **(E)** samples. The Venn diagrams show overlap between DT genes that were identified in female samples and in all samples (male and female). In the D1 samples, there was more overlap between female DT genes and DT genes in all samples than there was between male DT genes and DT genes in all samples. This raises the possibility that changes in the female rats were driving the large number of DT genes that we observed in the D1_AD40 Coc vs. AD1 Coc comparison.

The table shown in Fig. 4B shows male-female comparisons. The comparison in the top row tested how many genes were differentially translated between females and males in D1:Cre rats and A2a:Cre rats, when samples were combined across AD and cocaine/saline self-administration. In D1:Cre rats there were 66 upregulated and 18 downregulated genes in females relative to males. In A2a:Cre rats there were 138 upregulated and 32 downregulated genes in females relative to males. These results indicate greater sex-dependent regulation in the A2a-TRAP samples than the D1-TRAP samples. The rest of the table (rows 2-4) shows comparisons that tested how sex affected gene regulation within experimental group. The pattern of more sex-dependent translational regulation in A2a-TRAP samples than D1-TRAP samples described above held in all of the contrasts between males and females.

The tables in Fig. 4C shows comparisons testing how the experimental groups differed within each sex. In female rats (left), there were 118 upregulated and 159 downregulated genes in D1_AD40 Coc vs. D1_AD1 Coc and 19 upregulated and 73 downregulated genes in A2a_AD40 Coc vs. A2a_AD1 Coc samples. In male rats (right), there were 9 upregulated and 29 downregulated genes in D1_AD40 Coc vs. D1_AD1 Coc and 32 upregulated and 48 downregulated genes in A2a_AD40 Coc vs. A2a_AD1 Coc samples. The greater number of DT genes in female vs. male rats in the D1_AD40 Coc vs. D1_AD1 Coc comparison is notable. To test the extent to which differences in the female rats were driving the differences observed between cocaine AD40 vs. cocaine AD1 samples when the sexes were combined, we determined the number of genes that overlapped in the female D1_AD40 Coc vs. female D1_AD1 Coc contrast and the overall D1_AD40 Coc vs. D1_AD1 Coc contrast (Fig. 4D). The large degree of overlap between the female only comparison and the overall comparison (73 genes upregulated and 135 downregulated in both females and all rats), coupled with the much smaller overlap in the male only and overall comparison (5 genes upregulated and 17 downregulated in both males and all), suggests that the changes observed between D1_AD40 Coc vs. D1_AD1 Coc (seen in Fig. 3) could be driven largely by changes occurring in females (see Discussion for consideration of these data in light of equivalent incubation of craving in the sexes). There was far less overlap between the female A2a_AD40 Coc vs. female A2a_AD1 Coc contrast and the overall A2a_AD40 Coc vs. A2a_AD1 Coc contrast (Fig. 4E, left) than was observed in the D1-MSN. However, the degree of overlap between the male A2a_AD40 Coc vs. male A2a_AD1 Coc contrast and the overall A2a_AD40 Coc vs. A2a_AD1 Coc contrast (Fig. 4E, right) was similar to that observed in the D1-MSN.

### Pathways identified by bioinformatic analysis

DT genes in the D1_AD40 Coc vs. D1_AD1 Coc comparison were used to query the enrichR and Panther databases to identify pathway and GO category enrichment, respectively. Table 1A shows pathways identified as significantly enriched by EnrichR ranked by p-value (left) and pathways selected as being of particular interest using p<0.1 as a cutoff (right). These latter were selected for their potential relevance to incubation of cocaine craving and/or motivated behavior. Table 1B shows enriched GO categories ranked by p-value (left), and enriched GO terms of interest selected using p<0.1 as a cutoff (right). DT genes from the A2a_AD40 Coc vs. A2a_AD1 Coc comparison were also used to query these databases. The results were overwhelmingly non-significant; only one pathway in one library had an adjusted p-value<0.05 - the Unfolded Protein Response from the EnrichR msigDB2020 library.

The list of pathways (Table 1A, left) and GO categories (Table 1B, left) ranked by p-value are notable for the prevalence of immune response-related terms supporting neuroinflammation as a process occurring over protracted abstinence from cocaine self-administration. From the ‘Selected’ list, we would like to draw attention, in both analyses, to the presence of pathways and GO terms related to regulation of protein turnover (i.e., mTORC1 signaling, Parkin-Ubiquitin Proteasomal System, Translation Factors, and Proteasome pathways). The ubiquitin-proteasome system (UPS) [47] and protein translation [48] have been implicated in addiction, and specifically in the incubation of cocaine craving (see Discussion). A comprehensive list of all the GO categories and enriched pathways for the D1_AD40 Coc vs. D1_AD1 Coc, as well as the equivalent A2a comparison, can be found in Supplementary Material.

## Discussion

The goal of this study was to identify differentially translating (DT) genes in D1-MSN and A2a-MSN associated with incubation of cocaine craving. Three observations stand out: 1) For both D1-MSN and A2a-MSN, there were few DT genes between AD1 saline vs. AD1 cocaine groups. 2) In contrast, a pronounced divergence was observed between AD40 cocaine vs. AD1 cocaine groups, and this was far more robust in D1-MSN. 3) There are sex differences in translating mRNAs. Below, we discuss these observations, comparing our results to prior studies.

### Comparison to prior TRAP-seq studies in NAc after cocaine exposure

Relatively few differences between saline and cocaine rats were seen on AD1 (0 and 2 DT genes in D1-MSN and A2a-MSN, respectively). This may seem surprising given many reports that cocaine alters gene expression in MSN. However, our study is the first to specifically analyze translating mRNAs in D1-MSN and A2a-MSN, by combining TRAP-seq and transgenic rats, in the NAcC after cocaine self-administration. The only other NAc TRAP-seq study after cocaine self-administration (in male mice) did not distinguish D1-MSN vs. A2a-MSN – instead, mRNA from a neuronal ensemble, defined based on its activation during cocaine self-administration, were sequenced [49]. Another critical distinction is that [49] was performed in NAcS; incubation-related plasticity may differ in NAcS and NAcC [18,19,21,22]. Two other studies used TRAP and transgenic mice to study NAc D1-MSN and D2-MSN [32,50] but both used non-contingent cocaine (which produces different adaptations than cocaine self-administration [16]) and analyzed acute effects of cocaine. Overall, our results identify cocaine-associated populations of DT genes distinct from those studied previously.

### Robust differences in DT genes emerge in D1-MSN of cocaine rats after protracted abstinence

In contrast to the cocaine vs. saline AD1 comparison addressed above, there was substantial divergence in translating mRNAs between cocaine rats on AD1 (before incubation) and AD40 (after incubation), and this effect was far more robust in D1-MSN. Our finding of more DT genes on AD40 is consistent with two whole-genome expression studies that compared NAc at different abstinence times from cocaine self-administration, but neither of these distinguished D1-MSN vs. D2-MSN [51,52]. Briefly, one study obtained rat NAc tissue after a regimen shown to produce incubation of craving [53] and, using microarrays, found more differentially expressed genes (DEGs) (sal vs. coc) on AD10 vs. AD1 [51]. The other (RNAseq in mice) found ∼8 times more DEGs (sal vs. coc) on AD28 than AD1 after a cocaine self-administration regimen that did not appear to produce incubation of craving [52]. Also, two RNAseq studies compared mouse NAc at different abstinence times from cocaine self-administration but focused on effects of cocaine re-exposure [54,55], which is a very different biological condition from abstinence, and these did not distinguish D1-MSN vs. D2-MSN.

Our finding of more DT genes in D1-MSN vs. A2a-MSN is consistent with studies showing greater synaptic plasticity in NAc D1-MSN following cocaine self-administration [e.g., [56]], including the recent demonstration that CP-AMPAR upregulation during incubation occurs exclusively in D1-MSN [57]. However, this is the first demonstration of more robust plasticity after incubation in D1-MSN at the level of translating mRNAs. Recent studies performed single nucleus RNA-seq from NAc tissue obtained 1-h after acute or repeated experimenter-administered cocaine and also found a stronger transcriptional response in D1-MSN [45,58]. Furthermore, they classified distinct subpopulations of D1-MSN and D2-MSN and identified one population of D1-MSN as especially sensitive to cocaine.

### Sex differences in DT genes

Our analyses identify sex differences in translating mRNAs, both in saline animals and during incubation of cocaine craving. Notably: 1) When combining all groups and testing the main effect of sex (p-adj.<0.05), there were 222 DT genes. 2) Combining cocaine and saline rats and comparing males vs. females within each line, there was more sex dependent regulation in A2a-MSN (170 DT) than D1-MSN (84 DT) (Fig. 4B). 3) When analyzing each sex separately and comparing AD40 Coc vs. AD1 Coc, we saw similar numbers of DT mRNA in A2a males (80 DT) and females (92 DT), but many more DT mRNAs in D1 females (277 DT) than D1 males (38 DT) (Fig. 4C). Interestingly, previous work has shown similar cocaine incubation in both sexes, except for female rats in estrus on the day of the seeking test, who express more robust incubation than males or non-estrus females (which do not differ) [59–61]. Likewise, we found no significant sex differences in incubation or associated CP-AMPAR plasticity in NAcC [26] although we did not stage the estrous cycle in this prior study or the present one. However, given the short duration of estrus and thus the low likelihood of a female being in this stage, effects of estrus are unlikely to have dominated our data. In conclusion, both sexes incubate but they show divergence in DT genes associated with cocaine incubation. Similar results have been reported for incubation of morphine craving [62].

### Validation of selected DT genes

Four genes that showed differential translation in D1_AD40 Coc vs. D1_AD1 Coc rats (*Fosb and Epb41l3, upregulated; Kcna5 and Grm5, downregulated)* and are also linked to plasticity during cocaine abstinence (see below) were assessed with qRT-PCR to confirm the TRAP-seq results. For the first three, qRT-PCR showed significant changes in the same direction indicated by TRAP-seq, while *Grm5* trended in the predicted direction (Fig. 3I-L). The rationale for selection of these genes follows: 1) *Fosb:* Alternative splicing of *Fosb* mRNA yields ΔFosB, a transcription factor that is more stable than other family members and causes lasting changes in gene expression [63,64]. A role for ΔFosB in modulating cocaine-induced plasticity specifically in D1-MSN has previously been identified [65–68] and the role of ΔFosB in addiction broadly has been reviewed extensively [63,64]. In general, immediate early genes have an important role in the response of D1-MSN in the NAc to cocaine [45,58,69]. Our data are remarkable in showing that even 40+ days after the last cocaine experience, translation of *Fosb* is still upregulated in D1-MSN. 2) *Epb41l3:* This gene codes for erythrocyte membrane protein 4.1B (also known as erythrocyte membrane protein band 4.1-like 3). Its protein product, and related isoforms, can regulate DA, mGlu1, and NMDA receptors [70–72], all of which are implicated in cocaine incubation [19,25,26,73–78]. 3) *Grm5:* Previous work has demonstrated reduced mGlu5 surface expression and association with Homer proteins in the NAcC during incubation of cocaine craving [25]. The functional significance of this is unclear but it may contribute, along with other factors [79], to the loss of mGlu5-CB1R LTD that occurs after cocaine incubation [80]. 4) *Kcna5:* This gene was selected based on implication of potassium channel-related genes in prior studies of cocaine self-administration (see next section).

### Prior studies of gene expression after cocaine self-administration and abstinence

The only other TRAP study after cocaine self-administration found a time-dependent reduction of *Kcnj2* (AD30 vs. AD1), which codes for the inwardly rectifying K^+^ channel Kir2.1, specifically in mouse NAcS ensembles activated by cocaine self-administration. The associated increase in excitability within these ensembles was linked to incubation of craving [49]. We did not detect differential translation of *Kcnj2* in either MSN subtype between AD1 Coc and AD40 Coc, although we did observe differential translation of other K^+^ channels and interacting proteins (Supplementary Table 1 and Fig. 3J). Interestingly, in the prior TRAP study, MSN outside the targeted ensemble showed reduced excitability after incubation [49]. Others have found that reduced excitability of unidentified MSN in NAc shell, at least in part attributable to the SK2 type of Ca^2+^-activated K^+^ channels, is necessary but not sufficient for incubation of cocaine craving [21] and references therein]. This study also found that in early withdrawal from cocaine self-administration there was hypo-excitability of MSN in shell but not core in mice [21]. These findings, along with others [18,19,21,22], suggest distinct plasticity cascades in core and shell subregions. This likely explains why we did not detect differential translation of *Kcnj2* or *Kcnn2* (which codes for the SK2 channel) in our NAcC samples.

In an RNAseq study comparing mouse NAc between cocaine and saline groups on AD1 or AD28 after cocaine self-administration (the regimen did not appear to produce incubation), mRNA coding for *Nr4a1*, a member of the orphan nuclear receptor family (aka Nur77), was increased on AD1 (relative to saline) which then regulated addiction-related target genes in a persistent manner via histone modifications, leading to a homeostatic reduction in cocaine seeking and cocaine CPP [52]. We did not observe changes in *Nr4a1* or its target genes, likely because our analysis was focused so differently (D1 vs. D2, core, actively translating mRNA, incubation of craving).

Transcriptomic studies in NAc have implicated nuclear retinoic acid (RAR) or retinoic X (RXR) receptor signaling in rewarding effects of cocaine [54,81] and in the incubation of craving and its interaction with environmental enrichment [82–85]. In addition to its canonical role as a transcriptional regulator, RA also acts through dendritic RARs to elicit a form of homeostatic plasticity dependent on increased dendritic translation of *Gria1* leading to synaptic insertion of CP-AMPARs [86,87], and our recent findings suggest that this cascade is recapitulated in NAcC during cocaine incubation [27,57,88]. However, changes in translation confined to dendrites would not necessarily be detected by TRAP studies using whole-cell pulldowns as starting material. Indeed, we did not detect differential translation of *Gria1* after incubation of cocaine craving.

The UPS is linked to the incubation of cocaine craving [47,89–91] so it is noteworthy that EnrichR analysis identified pathways associated with the UPS (Table 1A). Among UPS-related genes, we observed that *Ndfip1*, the gene that codes for Nedd4 family interacting protein-1, was downregulated in cocaine D1-MSN on AD40 vs. AD1. Nedd4 is an E3 ubiquitin ligase implicated in AMPAR regulation [92–94]. A reduction could contribute to the accumulation of synaptic CP-AMPARs. Other E3 ubiquitin ligases, SMURF1 [91] and TRIM3 [90], are involved in cocaine seeking after cocaine self-administration and an abstinence period.

Finally, there was a prominent presence of immune response and inflammation response pathways that were significantly altered in D1_AD40 Coc vs. D1_AD1 Coc groups. Neuroinflammation and synaptic remodeling pathways were found to be robustly altered in the NAc of brains from humans with opioid use disorder [95]. Links between inflammation and psychostimulant exposure have previously been reported in rodents [96,97] and humans [98] [see [99] for review]. However, we cannot rule out an immune response to virus infusion.

## Limitations of our study

A limitation of our data set is the lack of an AD40 saline group. This group, when compared to AD1 saline rats, would help identify changes in gene expression related to duration of post-surgical single housing. It is worth noting that the rats in the AD40 Coc and AD1 Coc/Sal groups were age-matched so that there were no differences in age between the AD40 and AD1 groups, and also, identifying DT genes associated with incubation is most simply achieved through comparing AD1 and AD40 cocaine groups. Another limitation of these studies is difficulty in determining biologically relevant effect sizes and p-values given that important changes could be occurring that are small in magnitude or occur in a subset of MSN [100], and thus do not reach statistical significance. Finally, validation of genes of interest is important but accomplishing this in a cell-type specific manner at the protein level is associated with technical challenges. Despite these limitations, our study provides a new window on MSN subtype-specific changes in protein translation that may contribute to incubation of cocaine craving.

## Supporting information

Gene ontology raw data

## Acknowledgements

We thank the OHSU Integrated Genomic Laboratory for RNA sequencing and Brett Davis and Lucia Carbone of the OHSU Knight Cardiovascular Institute Epigenetic Consortium for their help with RNA-seq data analysis. We also thank Dr. Alexander Nectow for generously providing the TRAP virus for initial experiments.

## Author contributions

ABK and MEW wrote the manuscript; ABK, MEW, and MG developed the experiments; ABK, JGH, and MMB conducted the experiments; JGH and MG provided input on TRAP procedures.

