## Supplementary Materials for "Changes in nucleus accumbens core translatome accompanying incubation of cocaine craving"

Alex B. Kawa[1], Joel G. Hashimoto[1,2], Madelyn M. Beutler[1], Marina Guizzetti[1,2], Marina E. Wolf[1]
[1] Department of Behavioral Neuroscience, Oregon Health & Science University, Portland, OR 97239; [2] VA Portland Health Care System, Portland, OR 97239

**Contents: Fig. S1-3, Supplementary Table 1, Supplementary Methods, and References for Supplementary Materials**

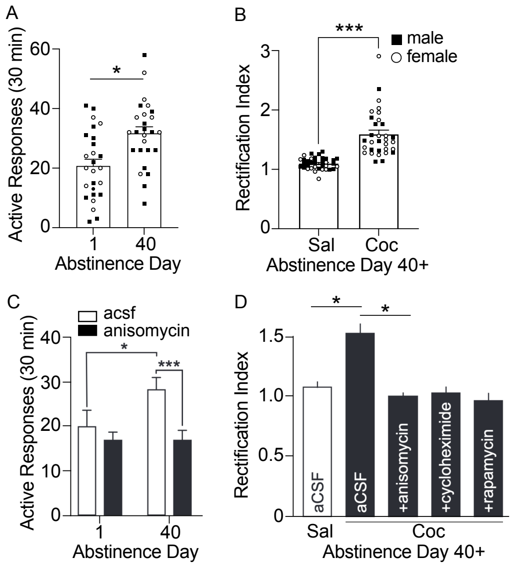


**Supplementary Fig. S1.** A summary of data supporting the role of protein translation in the incubation of cocaine craving (re-plotted from previous publications from the Wolf Lab). **A)** Rats make more responses at the previously active nose port on Abstinence Day 40 than Abstinence Day 1. This abstinence-dependent intensification of cue-induced drug seeking (incubation) has been observed in rats and humans across a number of drug classes. **B)** Whole cell patch clamp recordings from medium spiny neurons (MSN) in the nucleus accumbens core (NAcC) reveal a higher rectification index (RI) in MSN recorded from incubated rats than MSN recorded from rats that self-administered saline. An elevated RI indicates a greater contribution of $Ca^{2+}$-permeable AMPA receptors (CP-AMPARs) to synaptic transmission. **C)** Behavioral expression of incubation of craving requires ongoing protein translation. In this study, rats were treated with intra-NAc anisomycin to inhibit protein translation 30 minutes prior to a seeking test, and this treatment blocked expression of incubation. **D)** Whole cell patch clamp recordings from NAcC MSN. Cocaine incubated rats show an elevated RI relative to saline controls when aCSF is bath applied, while bath application of protein translation inhibitors normalizes the elevated RI present in MSN from incubated rats. These findings, together with others, demonstrate that ongoing protein translation is necessary for expression of incubation and maintenance of the upregulated CP-AMPARs upon which incubation depends. Data replotted from [1] (**A, B**), [2] (**C**), and [3] (**D**).

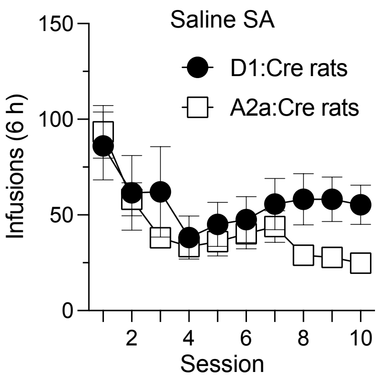


**Supplementary Fig. S2.** Saline self-administration data over 10 daily sessions in D1:Cre (n=8) and A2a:Cre (n=8) rats. Pokes in the active hole resulted in an intravenous infusion of saline paired with a light cue. D1:Cre and A2a:Cre rats did not differ in the number of infusions self-administered [effect of genotype, $p>0.05$].

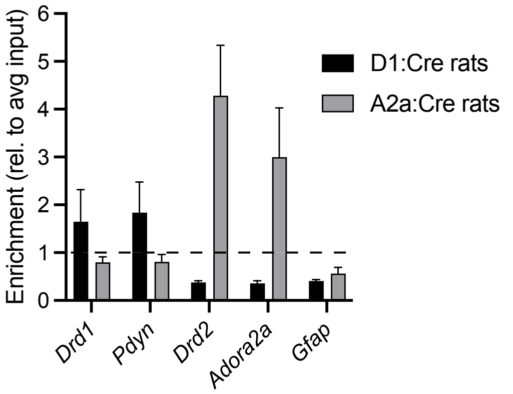


**Supplementary Fig. S3.** RT-PCR data from translating mRNA obtained from D1:Cre (n=2) and A2a:Cre (n=2) samples. The genes tested are markers of D1-MSN (*Drd1* and *Pdyn*), D2-MSN (*Drd2* and *Adora2a*) and astrocytes (*Gfap*). Relative to homogenate (input) mRNA (dashed line), the mRNA of D1:Cre rats was enriched for *Drd1* and *Pdyn* and de-enriched for *Drd2* and *Adora2a*. The mRNA of A2a:Cre rats was enriched for *Drd2* and *Adora2a* and de-enriched for *Drd1* and *Pdyn*. The mRNA from both cell types was de-enriched for *Gfap*, as expected.

**Potassium channels and interacting proteins**

| Gene | Change in D1-MSN on AD40-50 vs. AD1 (Coc rats) | Brief Description |
|---|---|---|
| *Kcna5* | down | potassium voltage-gated channel |
| *Kcnip1* | up | potassium voltage-gated channel interacting protein |
| **Gene** | **Change on AD40-50 vs. AD1 (Coc rats, D1 and A2a samples combined)** | **Brief Description** |
| *Kcnk4* | up | potassium two pore domain channel |
| *Kcnk3* | up | potassium two pore domain channel |
| *Kcnj14* | up | inwardly rectifying K+ channel |

**Supplementary Table 1**. Differentially translated mRNAs for potassium channels and interacting proteins identified by TRAP-seq analysis of NAc core on abstinence day (AD)1 versus AD40-50 following extended-access cocaine self-administration. See main text for discussion of functional significance.

### Supplementary Methods

Subjects

Procedures were approved by the Oregon Health & Science University Institutional Animal Care and Use Committee in accordance with the U.S. Public Health Service Guide for Care and Use of Laboratory Animals. D1:Cre and A2a:Cre lines were maintained by crossing with wild-type Long-Evans rats to maintain genetic diversity. All offspring were genotyped using established primers to ensure that only rats of the desired genotype were assigned to experimental groups. D1:Cre and A2a:Cre rats are available from the Rat Resource and Research Center (Columbia, MO; D1:Cre: RRRC#856, A2a:Cre: RRRC#857). All rats weighed at least 200 grams and were 8-11 weeks old before the start of the experiment. Rats were maintained on a reverse 12 h light/dark cycle with food and water available *ad libitum*.

RNAscope in situ hybridization

Brains (n=6 rats, 3 D1:Cre and 3 A2a:Cre) were flash frozen and stored at -80°C for approximately 2-3 weeks. Brains were sectioned on a cryostat at 16 μm and mounted on glass slides. Sections were fixed in 10% neutral buffered formalin at 4°C for 15 minutes and dehydrated through 50%, 70%, 100% and fresh 100% EtOH at room temperature (RT) for 5 min each. A hydrophobic barrier was drawn around each section. Hydrogen Peroxide was applied to each section for 10 min at RT. Slides were rinsed twice in water and then twice in 1x PBS (3 min each) at RT. Fluorescent probes (RNAScope, Advanced Cell Diagnostics, *iCre* [catalog 423321-C3], *Drd1a* [317031-C1], *Drd2* [315641-C1], and *Adora2a* [450471-C1]) were applied (2 h, 40°C) following a manufacturer-specified protocol (RNAscope Multiplex Fluorescent v2 protocol; Advanced Cell Diagnostics). DAPI was added to the slides before coverslipping (Prolong Gold, Thermo Fisher Scientific).

Intravenous Jugular Catheter Surgery

Rats underwent surgery to implant a silastic catheter (PlasticsOne, Roanoke, VA) into the right jugular vein, as described previously [1,4]. Briefly, for this surgery and others described below, rats were anesthetized using inhalable isoflurane (~5% for induction and then maintained at ~2%). Meloxicam (5 mg/kg, subcutaneous, Covetrus, Portland, ME) was administered during the surgical procedure and once daily for 48 h following completion of the surgery. After anesthesia was achieved, the catheter was inserted and secured into the jugular vein, and connected to a subcutaneous back mount, exiting the back in the mid-scapular region. Following completion of the surgery, rats were monitored until they were awake and active, and were singly-housed for the duration of the experiment. Intravenous catheters were flushed daily after surgery and through self-administration training with cefazolin (0.2 mL of 0.1 g/mL in sterile 0.9% saline) to prevent infection and maintain catheter patency. After 5-7 days of recovery, rats underwent 10 days of self-administration training.

Intracranial Virus Infusion

All of the TRAP-seq rats received intracranial surgery to infuse 1 µL of TRAP virus (EF1a-AAV5-FLEX-GFPL10a) into the nucleus accumbens (NAc) core (NAcC) (AP: +1.7; ML: +/- 2.4; DV: -7.0 at a 6° angle). TRAP virus was initially provided by Dr. Alexander R. Nectow (Columbia University) and then purchased from Addgene (catalog 98747-AAV5). Rats were anesthetized using isoflurane as described for the catheter surgeries. Virus was injected bilaterally at a rate of 250 nL/min, and the injector tip was left in place for an additional 10 min to allow for diffusion of the virus. The incision was sutured after injection and rats were returned to their home cages. Preliminary experiments indicated that virus expression and mRNA obtained through immunoprecipitation (IP) were maximal after 4-5 weeks of virus expression. Thus, the TRAP virus was expressed for 4-5 weeks and then rats were euthanized, NAcC punches were taken, and tissue prepared for TRAP IP (see below). Rats that were euthanized in early abstinence (Abstinence Day 1 (AD1)) had the virus infused ~2 weeks prior to the catheter surgery and rats that were euthanized in late abstinence (AD40) had the virus infused in the week following self-administration, allowing the virus to express for the same amount of time in both groups of rats (see Fig. 1A). Rat body weight and potential signs of illness were monitored daily until full recovery from surgery.

Extended-access self-administration

Rats were allowed to self-administer cocaine (0.5 mg/kg/infusion) or saline for 6 h/day for 10 days on a fixed ratio 1 schedule as previously described [1,5]. Cocaine HCl was obtained from the National Institute on Drug Abuse Drug Supply Program and dissolved in 0.9% saline. Briefly, we used operant chambers (MED Associates, Fairfax, VT) held within sound-attenuating cabinets and equipped with two nose-poke holes. Responding in the active hole resulted in an intravenous infusion of cocaine (~2.5 seconds) and the illumination of a cue light within the nose-poke hole for 4 sec. Nose pokes that were made during the 4-sec cue light presentation were recorded but had no consequences. Responding in the inactive hole was recorded but had no consequences. Rats were tested with methohexital sodium (~1 mg/kg, 10 mg/mL in sterile water, PAR Pharmaceuticals, Irvine, CA) to confirm catheter patency once prior to beginning self-administration and once after the last day of self-administration. If a rat's catheter was not patent at either checkpoint, or if the rat failed to discriminate between the active and inactive nose ports during self-administration, that rat was excluded from further testing. A total of 3 rats were excluded from the study for these reasons. Rats that self-administered saline were not required to discriminate between the active and inactive nose ports.

Translating RNA affinity purification

TRAP was carried out on all 63 brains from the rats that completed the cocaine or saline self-administration procedure, as previously described [6-8]. AD1 rats were taken from their home cages and euthanized ~18 h after the last self-administration session. AD40-50 rat were taken from their home cages and euthanized at approximately the same time of day. Punches were taken using a 2 mm punch tool (actual diameter of each punch is about 1.6 mm) centered over the NAcC were obtained from 2 mm think coronal slices. Next, homogenization buffer (HB) was used as described [8], with heparin omitted. Lysed tissue in HB were centrifuged at 10,000 g for 10 min at 4°C to remove insoluble debris, and the supernatant was used for all subsequent steps. Input samples, equivalent to 12.5% of the initial HB volume, were saved from the supernatant prior to the immunoprecipitation, and RNA was extracted at the same time as TRAP samples. 6.25 μg of two GFP antibodies (bioreactor supernatant, clones 19C8 and 19F7 from the Memorial Sloan-Kettering Institute Monoclonal Antibody Facility) was added to each sample for 4 h at 4°C under gentle end-over-end rotation. Samples were transferred to tubes containing 25 μl of Pierce Protein A/G Magnetic Beads (Thermo Fisher Scientific; 88803; lot VF298063) and incubated overnight at 4°C with end-over-end rotation. The following day, beads were washed in high-salt buffer as described [8], and RNA was isolated from the beads and input samples using TRIzol (Thermo Fisher Scientific, 15596018) and the Direct-zol RNA Microprep Kit (Zymo Research, R2062). RNA concentration was determined using the Quant-iT RiboGreen RNA Assay (Thermo Fisher Scientific, R11490) using a CLARIOstar Plus plate reader (BMG Labtech). TRAP IP controls with either no antibody or using wild-type tissue resulted in no measurable RNA based on the RiboGreen assay and only trace amounts of RNA detected using qRT-PCR.

Validation of Cre recombination, TRAP IP, and RNA-Seq by qRT-PCR

Confirmation of the TRAP IP protocol and of specific Cre recombination in D1- or A2a-MSN was carried out with qRT-PCR as described previously [7,9]. Primers were used with 5 ng of RNA and the Luna Universal One-Step RT-qPCR Kit (NEB) with SYBR Green detection on the CFX96 Real-Time System (Bio-Rad). Cycle threshold data were normalized to total RNA using RiboGreen

(ThermoFisher) and expressed as $log_2$ transformed data relative to the input control samples, with significance determined by the Student's *t*-test. Primers for rat: *Drd1* (Forward: 5'-AAGTCCCCGGAAGTGTGTTC-3'; Reverse: 5'-CAGGTGTCGAAACCGGATG-3'), *Drd2* (Forward: 5'-GCAGTCGAGCTTTCAGAGCC-3'; Reverse: 5'-TCTGCGGCTCATCGTCTTAAG-3'), *Adora2a* (Forward: 5'-ATTCCACTCCGGTACAATGG-3'; Reverse: 5'-AGTTGTTCCAGCCCAGCAT-3'), *Pdyn* (Forward: 5'-GCCTAGGAGTGGAGTGTTCG-3'; Reverse: 5'-GGGATAGAGCAGTTGGGCTG-3'), *Gfap* (Forward: 5'-CCTTGCGCGGCACGAACGAG-3'; Reverse: 5'-CCGAGCGAGTGCCTCCTGGT-3'). Confirmation of RNA-seq via qRT-PCR was carried out as described above with the following primers: *Epb41l3* (Forward: 5'- GGATGGCTCGGAGATCCTCA-3'; Reverse: 5'-CTTCGTTTCCAGTTTCTGCACC-3'), *Fosb* (Forward: 5'- AATGCAGAAACCGTCGGAGG-3'; Reverse: 5'- AGCCGTCTTCCTTAGCGGAT-3'), *Grm5* (Forward: 5'-CTGGGTTGCATGTTTGTCCC-3'; Reverse: 5'- TTTTCCGTTGGAGCTTAGGGT -3'), and *Kcna5* (Forward: 5'- GGCTGGAGAGGAGACCTACG-3'; Reverse: 5'- CCTCCTCCTCTGAGGGTCAT-3'). RNA-seq confirmation by qRT-PCR was analyzed for significant effects of abstinence day, sex, and sex by abstinence day interaction by two-way ANOVA followed by one-tailed t-tests with sexes collapsed based on the expected direction of regulation based on the results of the RNA-seq analysis.

<u>RNA-Seq and analysis</u>

RNA quality was assessed using an Agilent Bioanalyzer, with all samples having a RIN of >7 (range 7.1–9.2). From the 63 samples that underwent TRAP IP, we selected 48 (8/group) for sequencing. RNA-seq libraries were profiled on a 4200 Tapestation (Agilent) and quantified by real-time PCR using a commercial kit (Kapa Biosystems/Roche) on a StepOnePlus Real-Time PCR Workstation (ABI/Thermo). Libraries were then sequenced on a NovaSeq 6000 (Illumina) by the OHSU Integrated Genomic Laboratory (IGL). Fastq files were assembled from the base call files using bcl2fastq (Illumina). The fastq files were trimmed with Trimmomatic [10] using the built-in filters for Illumina adapters. After trimming, sequence files were aligned with the STAR aligner [11]. The reference genome was Rattus norvegicus Rnor_6.0, downloaded with annotations from Ensembl. Following the alignment, the SAM files were converted to BAM format using SAMtools [12]. Raw read count data were analyzed using the R package DESeq2 [13]. The significance of RNA-Seq data was determined using DESeq2 FDR-adjusted Wald test with adjusted p-values of <0.05. All processed and raw sequencing reads are publicly accessible at the Gene Expression Omnibus (GEO; https://www.ncbi.nlm.nih.gov/geo/) using accession number GSE263646. Genes with less than a total of 10 reads across the 48 samples were excluded from the analysis. Clustering and principal component analysis were conducted on all 48 samples.

Bioinformatics analysis

We limited the analysis to the following databases to focus our results on GO categories and known gene pathways: “GO_Molecular_Function_2021,” “GO_Cellular_Component_2021,” “GO_Biological_Process_2021,” “BioPlanet_2019,” “Elsevier_Pathway_Collection,” “KEGG_2021_Human,” “MSigDB_Hallmark_2020,” and “WikiPathway_2021_Human.” Tables 1A and 1B show GO categories and Gene Pathways ranked by p-value and also categories and pathways that we selected as relevant from a collection with unadjusted p-values <0.1.

Statistical analyses

Linear mixed models (LMM) were used for repeated measure self-administration data. The best-fitting model of the repeated measure covariance was determined by the lower Akaike information criterion score [14]. For the LMM analysis, depending on the model selected, the degrees of freedom may have been adjusted to a non-integer value. For behavioral data, SPSS (v29, IBM) and GraphPad Prism 9.2 were used for statistics and figures, respectively.


**Formatted:** Font: (Default) Arial
